# The integration of network biology and pharmacophore modeling suggests repurposing Clindamycin as an inhibitor of pyroptosis via Caspase-1 blockage in tumor-associated macrophages

**DOI:** 10.1101/2024.01.18.576201

**Authors:** Adrian Weich, Cindy Flamann, Johannes Berges, Krishna Pal Singh, David Chambers, Xin Lai, Olaf Wolkenhauer, Carola Berking, Gerhard Krönke, Shailendra Gupta, Heiko Bruns, Julio Vera

## Abstract

**Background:** Uveal melanoma (UM) is a highly malignant intraocular tumor with a poor prognosis and response to therapy, including immune checkpoint inhibitors (ICIs), after the onset of liver metastasis. The metastatic microenvironment contains high levels of tumor-associated macrophages (TAMs) that correlate positively with a worse patient prognosis. We hypothesized that one could increase the efficacy of ICIs in UM metastases by immunomodulating UM-associated macrophages.

**Methods:** To identify potential targets for the immunomodulation, we created a network-based representation of the biology of TAMs and employed (bulk and single-cell) differential gene expression analysis to obtain a regulatory core of UM macrophages-associated genes. We utilized selected targets for pharmacophore-based virtual screening against a library of FDA-approved chemical compounds, followed by refined flexible docking analysis. Finally, we ranked the interactions and selected one novel drug-target combination for *in vitro* validation.

**Results:** Based on the generated TAM-specific interaction network (3863 nodes, 9073 edges), we derived a UM macrophages-associated regulatory core (74 nodes, 286 edges). From the regulatory core genes, we selected eight potential targets for pharmacophore-based virtual screening (YBX1, GSTP1, NLRP3, ISG15, MYC, PTGS2, NFKB1, CASP1). Of 266 drug-target interactions screened, we identified the interaction between the antibiotic Clindamycin and Caspase-1 as a priority for experimental validation. Our *in vitro* validation experiments showed that Clindamycin specifically interferes with activated Caspase-1 and inhibits the secretion of IL-1β, IL-18, and lactate dehydrogenase (LDH) in macrophages after stimulation. Our results suggest that repurposed Clindamycin could reduce pyroptosis in TAMs, a pro-inflammatory form of programmed immune cell death favouring tumor progression.

**Conclusion:** We were able to predict a novel Clindamycin-Caspase-1 interaction that effectively blocks Caspase-1-mediated inflammasome activity and pyroptosis in vitro in macrophages. This interaction is a promising clinical immunomodulator of the tumor microenvironment for improving ICI responsivenss. This work demonstrates the power of combining network-based transcriptomic analysis with pharmacophore-guided screening for *de novo* drug-target repurposing.

**Graphical Abstract:** 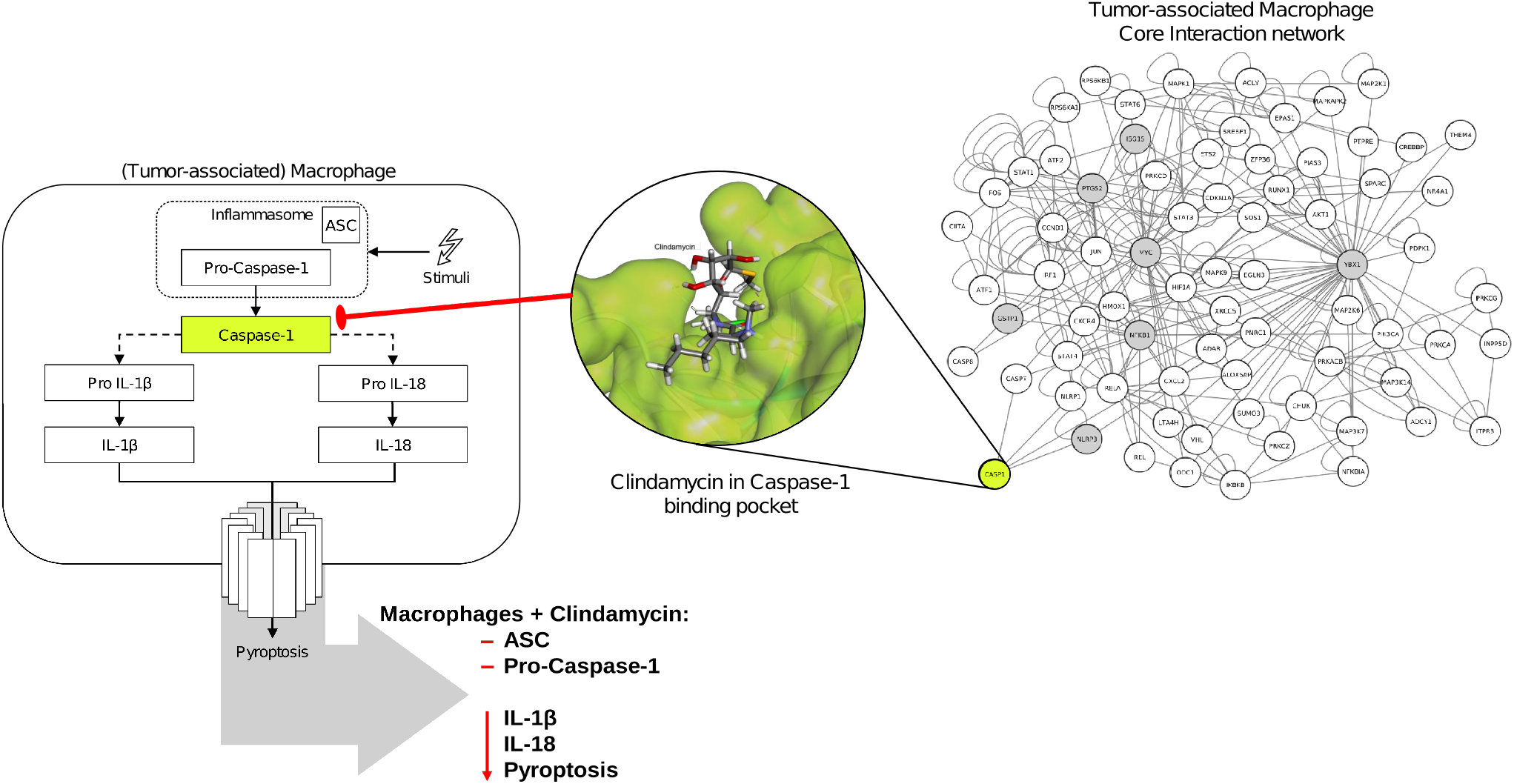

## Introduction

### The potential of immunomodulation of tumor-associated macrophages in uveal melanoma

Macrophages are among the most prevalent tumor-infiltrating immune cells. They have been observed to alter the effects of immune-checkpoint inhibition (ICI) therapy [1,2]. Uveal melanoma (UM), the most common ocular malignancy in adults, has a poor prognosis due to its liver metastases being extremely refractory to any therapy, including combined ICI therapies [3]. Since tumor-associated macrophages (TAMs) alter ICI responsiveness in other tumor entities, they may exert a similar effect in the metastatic UM. Moreover, TAMs in UM promote disease progression, and high levels of TAMs positively correlate with poorer prognosis and shorter survival of patients [4,5]. Thus, we hypothesize that immunomodulation of TAMs in UM can be employed to remodel the tumor microenvironment and help increase ICI responsiveness in UM. Recent findings support this hypothesis by showing that IL-1β, a central effector molecule following macrophage activation, drives pancreatic ductal adenocarcinoma growth, and its inhibition lowers inflammatory levels [6]. To explore this hypothesis, we developed a computational model of TAMs that can systematically identify important TAM regulatory factors exerting tumor-critical functions. This approach can potentially find therapeutic targets for the immunomodulation of TAMs.

### Computational Drug Repurposing

We propose drug repurposing, i.e., the use of existing drugs for a clinical purpose different from what they were initially approved for, to therapeutically influence the identified targets. With drug repurposing, one can utilize prior information about the biodistribution and toxicity of existing drugs to speed up their re-utilization and reduce the time from discovery to clinical approval [7,8]. Drug repurposing is also aligned with the procedure followed by molecular tumor boards with patients not responding to standard-of-care therapies. Traditionally, drug repurposing is often investigated utilizing systematic *in vitro* screening of drugs [9]. Many of the successfully repurposed drugs have been used on their original molecular target but for a different clinical condition [8]. However, one can repurpose drugs to new molecular targets utilizing computational biology. Goody and co-workers, for example, combined docking simulation-based screening of an FDA-approved molecule library and *in vitro* experiments to repurpose Argatroban to interfere with the interaction between metastasis-associated protein 1 (MTA1) and the cancer transcription factor E2F1, a molecular target unknown for this drug [10].

The patient -omics data analysis can speed up drug repurposing [11]. Cancer proteins are not isolated but belong to large gene and protein networks. Thus, one can combine -omics data and network biology algorithms to select protein targets for drug repurposing [12,13]. Here, we present an integrative computational workflow that combines transcriptomic data and network-driven selection of proteins as molecular targets with pharmacophore modelling of an FDA-approved drug library to repurpose drugs for them. We deployed the workflow using the targeting of TAMs in UM as a case study, although the methodology and key results are not limited to this tumor entity.

Further, we utilized *in vitro* experiments to validate the predictions. This enabled us to discover a novel interaction between the antibiotic Clindamycin and activated Caspase-1, which harbors the potential to inhibit the secretion of pro-inflammatory cytokines like IL-1β to the macrophage-surrounding environment, thereby preventing pyroptosis, a pro-inflammatory form of programmed immune cell death. [14].

## Materials and Methods

### Workflow

To repurpose drugs to target tumor-associated macrophages (TAMs), we implemented the following workflow (Figure 1):

1. **TAM network construction:** We collected bulk RNA sequencing data and signaling path-way data from public repositories and the literature. The latter was used to construct a regulatory network of biological interactions, while the former was used to achieve TAM-specificity via projecting the gene expression data onto the respective network nodes.
2. **Core network detection:** We extracted regulatory motifs from the network and ranked them based on their potential importance for the TAMs. Scoring parameters were network features (node degree, betweenness centrality) and differential expression data derived from publicly available single-cell RNA-Seq (scRNA-Seq) of UM-associated macrophages and healthy control macrophages.
3. **Docking simulations:** After selection of potential targets from the core network, we generated pharmacophore models of the respective proteins and performed virtual screening of FDA-approved drugs. For a selection of high-affinity candidates, we applied refined flexible docking with their potentially binding chemical compounds.
4. **Validation experiments:** We performed *in vitro* validation experiments using macrophages to validate one selected completely novel drug-target interaction.

**Figure 1.**
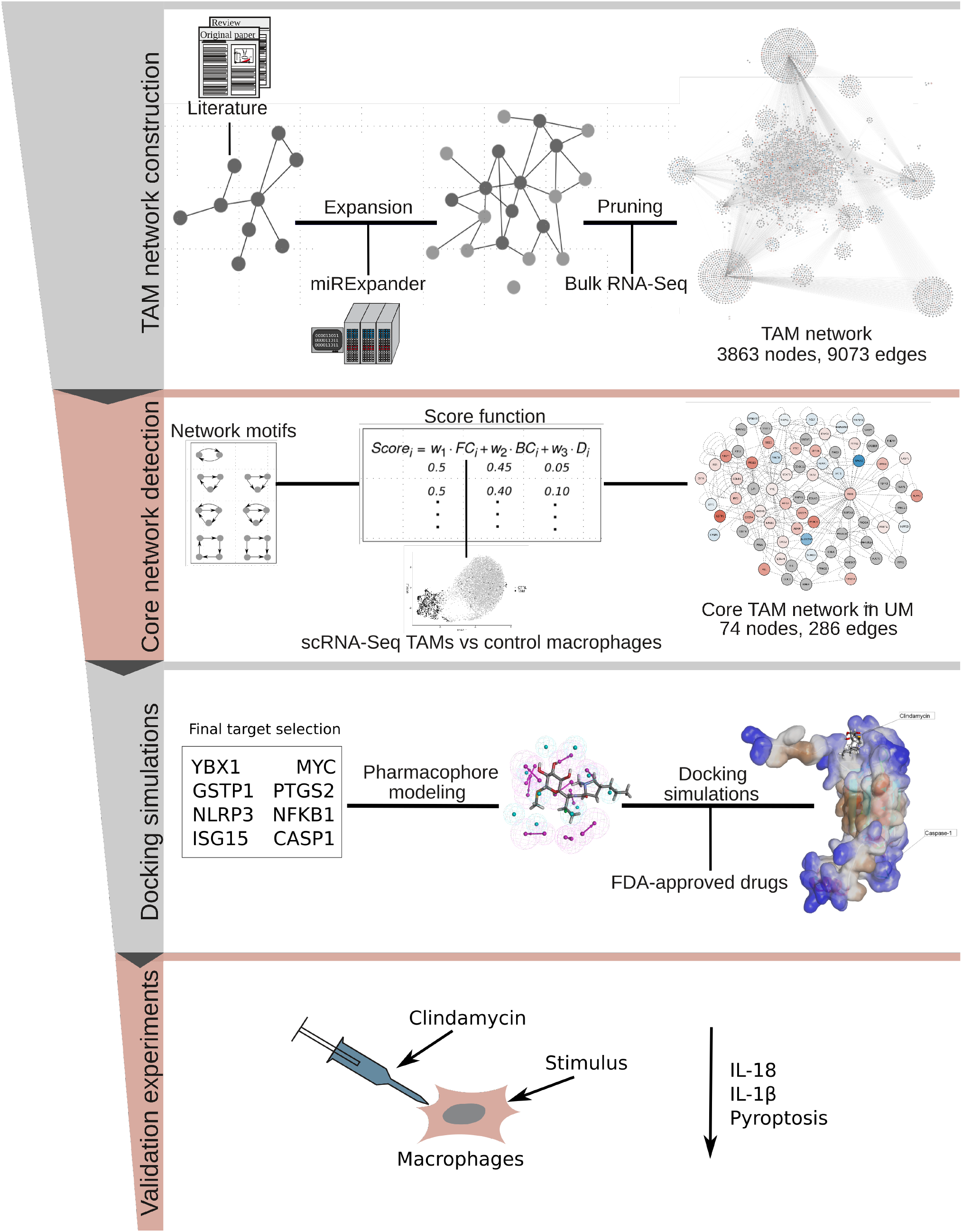
Workflow designed to detect, select, and test molecular targets and drugs for repurposing.

In the following, one can find a detailed explanation of the individual steps in the workflow.

### Data Collection

We obtained the different sequencing datasets from the GEO database. The data used for the network specification consisted of 12 bulk RNA sequencing samples (GSE117970) of macrophages associated to breast or endometrial cancer [15]. The data used for the differential expression analysis consisted of single-cell RNA sequencing results of 8 primary and 3 metastatic uveal melanoma samples (GSE139829, [16] and a collection of samples from healthy joint macrophages (GSE134691, [17].

### Differential Expression

Following the analysis workflow of the original publication, we combined 8 primary and 3 metastatic tumor samples in R (4.05) and aggregated them into a Seurat object with the “min.features” option set to 120 (Seurat V3) [18,19]. To extract only the TAMs from the Seurat object, we used the “Subset” function with the macrophage identifiers CD68, CD163, and CD14, each showing an expression greater than 1. These high criteria were used to reduce the false positive cells in the data, thereby assuring that the cells selected were true macrophages and avoiding contamination from other cell types. We then combined the TAMs subset with the healthy joint macrophages in the same Seurat object using the “merge” function. We set the identities of the TAMs and the healthy joint macrophages to “ident.1” and “ident.2”, respectively. We considered cells with more than 25% of all features being mitochondrial genes contamination and discarded them. After scaling, normalization, and principal component analysis using standard procedures, we performed batch correction using harmony [20]. The impact of the batch correction is illustrated in Figure S1. For the sets of cells identified in the Seurat object, we performed the differential expression analysis utilizing the “FindMarkers” function. We included only genes that were at least expressed by some cells of both conditions with “pct.1 > 0” and “pct.2 > 0” (Pct.1: percentage of cells in group 1, TAMs, expressing a specific gene). For these genes, we further selected the genes that have an adjusted p-value < 0.05 (Bonferroni correction). We exported the significant genes and their average log_2_ fold-change (log_2_FC) values for their use in the core network extraction. The plots were generated using Seurat’s “DimPlot”, “FeaturePlot”, and “DoHeatmap” functions.

### Network construction

The TAM network is based on the previously published macrophage network by Wentker *et. al* [21]. As this network only displays the M1-like polarization type of macrophages and TAMs are known to play a bilateral role in cancer, we extended the network with M2-like macrophage behavior [22–24]. To this end, we manually queried the NCBI’s PubMed archive for terms concerning the M2-like macrophage phenotype, including “M2 macrophage polarization”, “alternative activation of macrophages”, and “anti-inflammatory macrophages”. We also browsed the literature for pathways, proteins, genes, with a focus on cytokine production or transcription factor regulation. This information was added to the existing macrophage map using CellDesigner (v4.4.2) [25,26], and each new interaction was annotated utilizing CellDesigner’s MIRIAM [27]. We separately annotated all factors involved in the interactions: genes were annotated with Ensembl IDs [28], proteins with UniProt IDs [29], microRNAs with miRbase IDs [30], and simple molecules and ions with ChEBI IDs [31]. We used IDs from either mouse or human depending on the organism described in the corresponding literature. The two organism-specifications were later collapsed into human-only by using the biomaRt package (version 2.56.0).

Next, we extended the network automatically with information taken from miRTARBase (version 6.1) [32], miRecords (version 4.5) [33], HTRIdb (version 1) [34], and TRANSFAC (version 2015.1) [35] using an inhouse tool named miRNExpander (https://github.com/marteber/miRNexpander). To this end, we transformed the network into a Graph Modelling Language (GML) and continued working with the expanded network using Cytoscape (v3.8.0) [36].

We specified the expanded macrophage network to a TAM network by pruning it with RNA sequencing data from 12 samples, derived from breast and endometrial cancer associated macrophages (GSE117970). To this end, we combined the RNA-Seq data in R (4.0.5) and transformed the counts to transcripts per million (TPM) using Ensembl transcriptome as transcript-length reference (version GRCh37.87). We calculated the average TPM value of a gene and added it to the expanded network. The restrictions for the preservation of a node were set to an average TPM of at least 10 and a node degree of at least one. We saved the pruned network as a Cytoscape file and exported a list of its nodes for its perusal. We added to the network the significant log2FC values derived from the single-cell uveal melanoma TAMs data. The obtained network can be browsed and downloaded from www.vcells.net/TAM.

### Gene set enrichment analysis

We conducted gene set enrichment analysis (GSEA) using EnrichR [37] with the Mammalian Phenotype Ontology database [38] and the genes from the differential expression analysis belonging to the TAM network. The resulting tabular data was visualized in R using ggplot2 and ComplexHeatmap [39].

### Topological Features and Motif detection

We calculated the networks topology features using the built-in Cytoscape “Analyzer” [40]. Two network topological features were especially interesting: the node degree or number of node interactions, and the betweenness centrality, which indicates how many shortest pathways include the node considered. We added these metrics to the network annotation. Further, we queried the TAM network for regulatory motifs using the Cytoscape app “NetMatchStar” [41]. We decided to include 2-edges-2-nodes feedback loops, 3-edges-3-nodes feedback loops, 3-edges-3-nodes feedforward loops, 4-edges-3-nodes feedback loops, 4-edges-3-nodes feedforward loops, 4-edges-4-nodes feedback loops, and 4-edges-4-nodes feedforward loops. The same strategy was used to identify network motifs in our previous publication [42].

### Motif ranking

To detect the most important nodes and their interactions, we calculated a weighted ranking score of the identified motifs with the following equation:

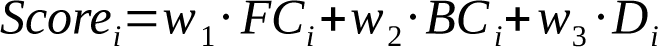

The score is based on the method used in Khan *et al.* [43]. For each motif *i*, the score is calculated with different weight settings for *w_1,_ w_2,_* and *w_3_* that define the importance of the three ranking factors. These factors are: a) *FC_i_* is the average log_2_FC expression in the scRNA-Seq from UM TAMs across the nodes forming the motif *i*; b) *BC_i_* is the average betweenness centrality of the motif *i*’s nodes; and c) *D_i_* is their average node degree. The weighting factors sum up to one and *w_1_* was fixed to 0.5 to prioritize motif expression when scoring motifs. We set the values of *w_2_* from 0.05 up to 0.45 in 0.05 iterative steps and the values of *w_3_* result from the calculation *w_3_*= 0.5 – *w_2_*. We calculated the motif scores of each motif *i* for each combination of weighting factors. Next, we pareto-optimized the different scores of the same motifs with the “psel” method using the R package rPref (version 1.3) [42,43].

### Core Extraction

We considered the components of the top 100 highest scoring motifs to be the core nodes [42,43]. Next, we extracted the core nodes and their interactions from the TAM network to create a core network, which can be browsed and downloaded from www.vcells.net/TAM

### Target Selection

We used a Min-Max-normalization metric to give us an idea about the relevance of each node in the core network:

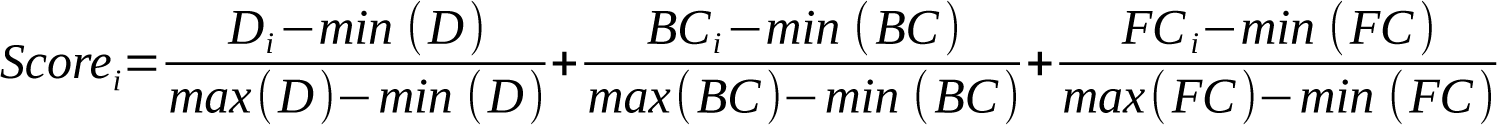

The score is based on the degree (D), betweenness centrality (BC), and differential expression (FC) of each node. We derived the topological features, namely degree and betweenness centrality from the core network, whereas we preserved the differential expression values from the TAM network. We used the ranking table to select 8 targets for pharmacophore modeling while already accounting for experimental suitability.

### Pharmacophore modeling and *in-silico* screening of drug library

We retrieved the 3D structures of the selected protein targets from the RCSB protein database (www.rcsb.org/pdb; PDB ID: 3QF2, 5X79, 1IBC, 3GUT, 6LMR, 3SDL, 5F1A; MYC via homology model). To each of them, we applied standard protein-preparation protocols of the Biovia Discovery Studio 2022 (DS 2022) to prepare them for pharmacophore generation. In this method, the features present in the active site of a protein act as a potential chemical fingerprint for drug screening. We used the ‘Edit and Cluster Features Tool’ of DS 2022 to generate the pharmacophore features from each active site of the proteins, including features like “Hydrogen Bond Donors and Acceptors” and “Hydrophobic”. We considered the excluded volume constraints to the best-selected pharmacophore model to highlight potentially forbidden sites for the drug molecules during the screening process. For the pharmacophore model screening, we utilized FDA-approved drugs in the Zinc15 database [44]. All the screened drugs were arranged in decreasing order of their FIT score, which represents how accurately a drug fits into the binding site. For each of the target proteins, we considered drugs that have a FIT value of more than 1. Afterwards, we searched for commonly screened drugs that could serve as potential targets for multiple proteins.

### Molecular docking

To further refine the prediction of the most promising drugs interacting with CASP1, YBX1, ISG15, and PTGS2, we performed a flexible docking on the binding site of the proteins. To this end, we extracted the binding site of the proteins from the experimental literature [45–48] and performed the flexible docking using the CDOCKER program of DS 2022. We generated 10 conformations for each of the drug–protein target combinations, which were ranked based on CDOCKER-estimated energy.

### Preparation of macrophages

We isolated human peripheral blood mononuclear cells (PBMCs) from freshly drawn peripheral blood of healthy donors (University Hospital of Erlangen, Department of Transfusion Medicine and Haemostaseology, GER) by density gradient centrifugation using human Pancoll (1.077 g/ml) (PAN™ Biotech, Aidenbach, GER) and a subsequent buffy coat purification. To generate macrophages, we isolated monocytes by adherence to polystyrene in CELLSTAR® cell culture flasks (Greiner Bio-One, Kremsmünster, AUT) and cultured in the presence of Leuco-max® GM-CSF (500 U/µl) (Novartis Pharma, Nürnberg, GER). After 6-7 d of culture, macrophages were detached with EDTA (1 mM) (Sigma-Aldrich®, München, GER).

### ELISA

We examined cell culture supernatants, serum levels for human IL-1β and IL-18 with ELISA kits from R&D Systems® (Minneapolis, USA) according to the manufacturer’s instructions.

### LDH release assay

We plated macrophages in 96-well culture at a concentration of 5 × 10^4^ cells/well and pretreated them with or without lipopolysaccharides (LPS, 100ng/ml) for 24 hours. Subsequently, we treated macrophages with Nigericin (10µM) in the presence or absence of Clindamycin (10µg/ml) overnight. LDH released in the supernatant was detected using a cytotoxicity detection kit (Roche) according to the manufacturer’s instructions. We used data on detected LDH to calculate the pyroptotic rate of treated macrophages based on the following equation: [(experimental release − spontaneous release)/(maximum release − spontaneous release)] × 100, where spontaneous release is from the cytoplasm of untreated macrophages, and maximum release is that obtained by lysis of macrophages with a solution of 0.1% Triton X-100.

### FLICA® 660 Caspase-1 assay

We detected Caspase-1 activity using the FLICA® 660 Caspase-1 assay kit from ImmunoChemistry Technologies (Bloomington, USA) according to the manufacturer’s instructions. We seeded macrophages at 1× 10^6^/ml in polystyrene Falcon® round bottom tubes (Corning® LifeSciences, Corning, USA) for flow cytometry. Cells were LPS-primed (100 ng/ml, 24 h) and overnight-incubated with 10 µM Nigericin in the presence or absence of Clindamycin (10µg/ml or 25µg/ml). We washed the cells with PBS and incubated with the FLICA® 660-YVAD-fmk reagent (1:150, 30 min) at 37 °C and 5 % CO2. As assessed by flow cytometry, Caspase-1 activation was defined as increase in red fluorescence.

### Western blot analysis

We seeded macrophages at 2× 10^6^/ml in polystyrene Falcon® 24 well plates (Corning® LifeSciences, Corning, USA), LPS-primed (1 µg/ml, 3 h) and overnight-incubated with 10 µM Nigericin in the presence or absence of Clindamycin (10µg/ml or 25µg/ml). We prepared cell lysates by direct lysis in 2 % (w/v) SDS lysis buffer (5 mM EDTA, 50 mM Tris/HCl, 150 mM NaCl, 2.2 % (wt/vol) SDS) supplemented with complete™ EDTA-free (Roche Diagnostics, Mannheim, GER) as protease inhibitor. We removed cell debris by centrifugation (21,382 xg, 15 min, 4 °C) and the concentration of total protein in cell extracts was determined using the Qubit® protein assay kit and the Qubit® 3.0 fluorometer (Thermo Fisher Scientific™). Cell culture supernatants were used purely. We suspended protein samples in 4× Laemmli sample buffer (278 mM Tris/HCl, 355 mM 2-mercaptoethanol, 0.02 % (wt/vol) bromophenol blue, 4.4 % (wt/vol) lithium dodecyl sulfate, 44.4 % (vol/vol) glycerol, pH (HCl) 6.8) (Bio-Rad Laboratories, München, GER) and boiled for 10 min at 95 °C. We separated the protein content of cell lysates, supernatants and the Precision Plus Protein™ WesternC™ standard (Bio-Rad Laboratories, München, GER) by SDS-PAGE (10 %, 15 %, 90 µg) and transferred onto nitrocellulose membranes (0.2 µm) (GE Healthcare Life Sciences, Chalfont St Giles, UK) using the semi-dry TransBlot® Turbo™ transfer system (Bio-Rad Laboratories, München, GER). We blocked membranes in 5 % (wt/vol) dried milk in TBS-T (100 mM Tris/HCl, 150 mM NaCl, 0.1 % (vol/vol) Tween®-20) for 1 h at room temperature. Membranes were overnight-incubated with primary antibodies diluted in 5 % (wt/vol) dried milk in TBS-T at 4 °C. Subsequently, we incubated membranes with the appropriate HRP-conjugated secondary antibody diluted in 5 % (wt/vol) dried milk in TBS-T for 1 h at room temperature. We detected proteins by chemiluminescence using the SuperSignal® ELISA femto maximum sensitivity substrate (Thermo Fisher Scientific™, Waltham, USA) according to the manufacturer’s instructions and the Amersham™ Imager 600 (GE Healthcare Life Sciences, Chalfont St Giles, UK). We stripped the membranes using the Restore™ western blot stripping buffer (Thermo Fisher Scientific™, Waltham, USA) before being re-examined. Primary antibodies used were β-actin (4967, Cell Signaling), ASC (AL177, AdipoGen/Biomol, 1:1000) (1:2.500) and Caspase-1 (clone: D7F10, Cell Signaling). Secondary HRP-conjugated antibodies used were anti-mouse IgG (7076, Cell Signaling) and anti-rabbit IgG (7074) (1:2,500) (Cell Signaling Technology®, Cambridge, UK).

## Results

### The combination of scRNA-Seq, bulk RNA-Seq, and network analysis generates a core network of potential molecular targets linked to immunomodulation and depletion of TAMs

We hypothesize that immunomodulation of TAMs in UM is a key for remodeling the tumor microenvironment which may ultimately help to increase ICI responsiveness in UM. To find molecular targets for immunomodulation in UM-associated macrophages we (a) constructed a comprehensive signaling and gene network reflecting the biology of TAMs, (b) integrated in the network nodes RNA-Seq data from TAMs and quantified their topological importance, and (c) used single-cell data and topological features to isolate a core network including the most connected and differentially expressed genes, and select from this core promising, druggable proteins (Figure 3A). Precisely:

**a) TAM network reconstruction**. To build a network representative of TAM biology, we expanded a previously published macrophage network by adding genes and pathways linked to the anti-inflammatory polarization of macrophages. This manual curation resulted in a network with 1318 nodes and 1014 edges. Next, we utilized a computational pipeline to further extend the network with molecules and interactions taken from protein-protein, transcriptional, and miRNA regulation databases. To remove irrelevant or poorly expressed genes from the network, we only conserved genes with an average TPM of at least 10 in RNA-Seq TAM data and a node degree of at least one in the network. This way, we obtained a fully-connected TAM network with 3863 nodes and 9073 edges (Figure 3B).
**b) Integration of scRNA-Seq data and topological features in the TAM network**. To quantify the importance of each node in the TAM network, we computed the topological features node degree and betweenness centrality. To fit the analysis as much as possible to our case study of TAMs in UM, we obtained scRNA-Seq datasets from primary and metastatic UM (GSE139829) and processed them utilizing Seurat [16]. To extract the TAMs from the Seurat object, we selected the individual cells that show an expression greater than one for the known macrophage surface markers CD68, CD163, and CD14. For the purpose of comparison and differential expression analysis, we utilized scRNA-Seq data sets from healthy macrophages (GSE134691). To allow the data integration, we applied scaling, normalization, batch correction, and performed differential expression analysis and p-value correction. Dimensionality reduction plots can be found in Figure S1. Following this approach, we extracted a group of TAMs consisting of 888 cells and combined it with a second group of healthy macrophages including 7542 cells (Figure S1C). The Seurat object measured 12172 features, of which 1671 were differentially expressed. We focused the analysis on the 3863 genes included in the TAM network as nodes, and identified 1367 genes that were differentially expressed with at least one cell per group expressing the respective gene feature, with 688 genes upregulated in the TAM group. A comparison of the transcriptomic patterns between the top 20 differentially expressed genes shows a clear upregulation in the TAM group of inflammatory proteins like IL-1β, NR4A2-3, TNFAIP3, or NLRP3, which are not similarly expressed in the healthy macrophage group (Figure 2A). TGFβ1, a gene known to play a role in inflammation and tissue regeneration, is expressed in both TAMs and healthy macrophages, albeit at different intensities [49,50]. These observations are in line with the generally upregulated phenotypes derived from the GSEA of the differentially expressed genes (n=1367) (Figure 3C). On one hand, we found dozens of enriched phenotypes related to abnormal physiology of macrophages including phagocytosis, chemotaxis, morphology, and differentiation (Figure 3C). On the other hand, we found several enriched phenotypes associated to the tumor microenvironment, including tumor necrosis factor secretion related with inflammation-associated carcinogenesis and tumor vascularization (Figure 3C). The distribution of all enriched phenotypes by the size of their gene sets can be found in Figure S2.
**c) Core network extraction and target selection**. We assumed that regulatory motifs like feedback and feedforward loops play a pivotal role in the (de)regulation of gene networks and isolated a core network composed of differentially expressed, highly connected and intertangled regulatory motifs. To this end, we first detected the 2-4 nodes feedback and feedforward loops contained in the network and obtained 9035 motifs (Table S1). We quantified their importance in terms of the topological features average node degree (*D_i_*) and betweenness centrality (*BC_i_*) of the nodes belonging to the motif. Also, we computed the average log_2_ fold change expression across the nodes forming each motif when comparing scRNA-Seq from TAMs and healthy macrophages (FC*_i_*). We combined these metrics into a computational score and used it to generate a core network containing the Pareto-optimized, top ranked network motifs (see Material and Methods). We obtained a core network with 74 nodes and 286 edges (Figure 3D). We generated a ranking of the most important nodes regarding differential expression between healthy and TAMs and their importance for the core network (see Material and Methods and Table S2). We combined the ranking information with foreseen experimental validation suitability and thereby selected eight potential drug targets among the high-scoring candidates for further investigations, namely: YBX1, MYC, GSTP1, PTGS2, NLRP3, NFKB1, ISG15, and CASP1. When we inspected the scRNA-Seq data, we found that these genes are to some extent expressed in both cell types, but with a higher intensity in the TAM group (Figure 2B). For instance, YBX1 shows a rather universal expression across all cells, whereas GSTP1 seems to be rather TAM-exclusively expressed.

**Figure 2.**
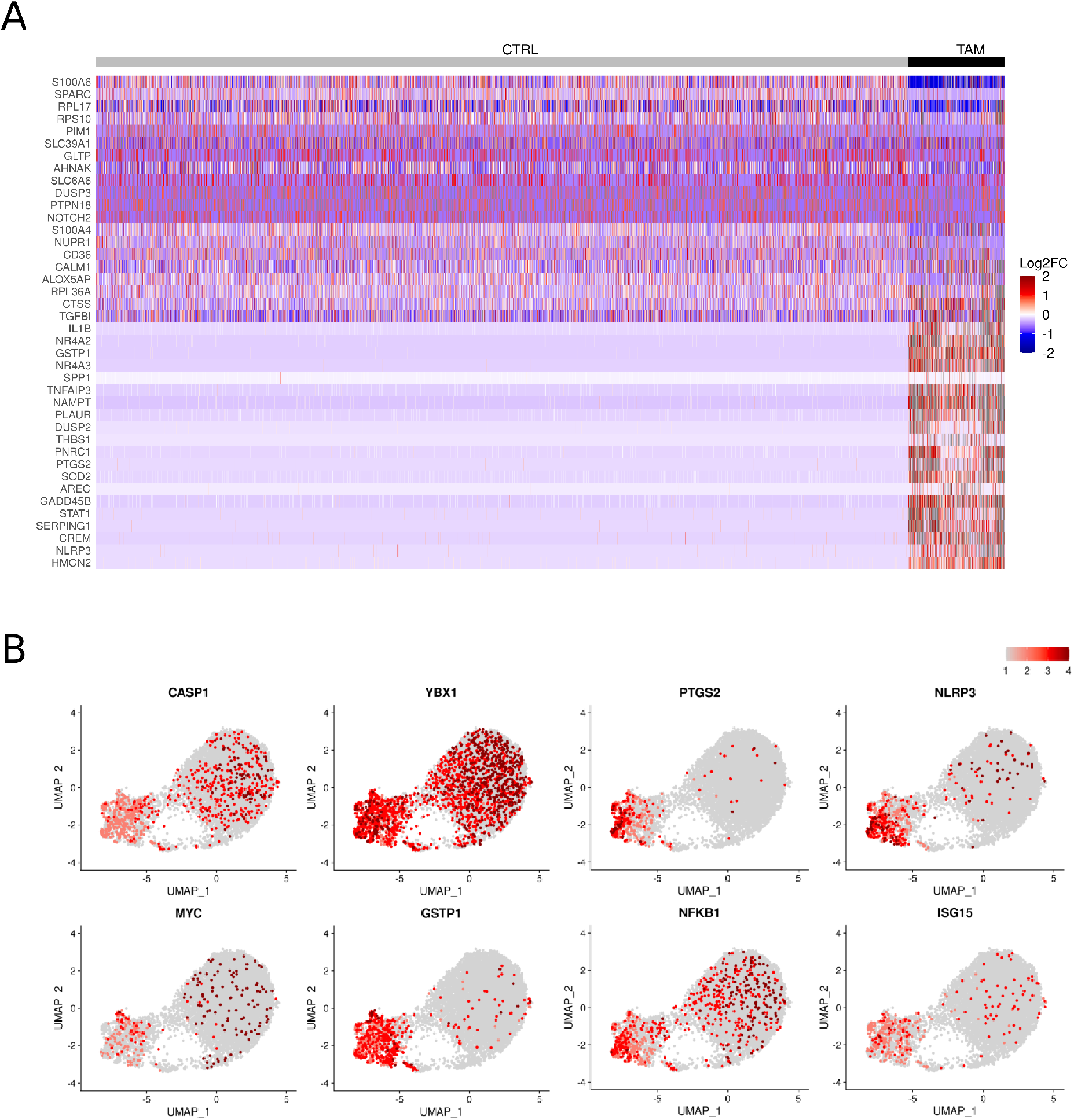
Single-cell expression analysis reveals differences between healthy macrophages and cancer-associated macrophages. **A) Heatmap of Top20 differentially expressed genes.** While the upregulated genes of the healthy joint macrophages (CTRL) group are also widely expressed in the TAM group, the top 20 genes upregulated in the TAM group show no such expression in the CTRL group. This indicates a specialized or generally more active phenotype of TAMs compared to the healthy subset. **B) Gene expression of the 8 targets selected for pharmacophore modeling.** PTGS2, NLRP3, and GSTP1 show a distinct TAM-exclusive expression. In contrast, YBX1, NFKB1, and CASP1 are active to at least some extent in TAM and CTRL cells. This hints towards a rather homeostatic role of the latter genes.

**Figure 3.**
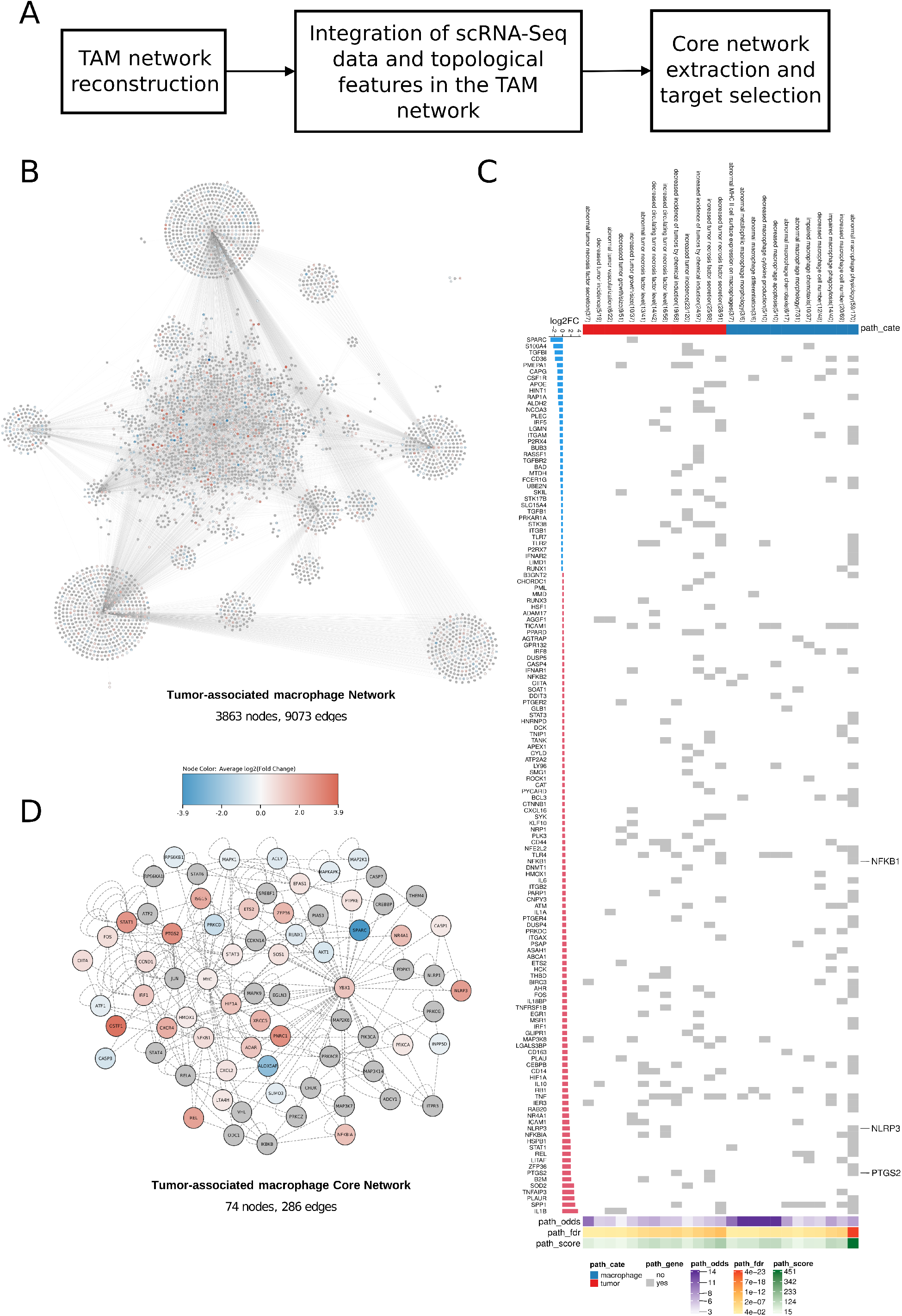
Identification of a core interaction network of tumor-associated macrophages. A) The workflow followed for core network extraction. B) TAM network. The TAM network consists of 3863 nodes and 9073 edges. It is based on literature, database knowledge about interactions of biological entities in macrophages, and bulk RNA-Seq expression data. **C) Gene-set enrichment analysis of differentially expressed genes with nodes in the network.** Input genes for the GSEA were derived from the overlap of differentially expressed genes from the single-cell data and the nodes in the TAM network (n=1367). Each grid in the heat map represents whether a gene is enriched in a phenotype. Only tumor- (red) and macrophage-related (blue) phenotypes were selected for visualization in the heatmap. In addition, we annotated each phenotype with its odds ratio (odds), adjusted p-values calculated using false discovery rate (FDR), and a combine score (odds x [-log10(FDR)]). The bar plot showed the corresponding log2FC of the enriched genes in the phenotypes. We highlighted NFKB1, NLRP3, and PTGS2 because they are among the eight targets selected for pharmacophore modeling. **D) TAM core network.** The TAM core network consists of 74 nodes and 286 edges. The nodes were colored according to their differential expression values (log2FC) derived from the single-cell data analysis. Nodes in grey showed no significant differential expression. When it comes to the 8 nodes selected for pharmacophore modelling experiments, each of them was at least slightly upregulated in the TAM group compared to the healthy macrophages.

### Pharmacophore modeling and docking simulations of FDA-approved chemical compounds suggests Clindamycin and other drugs for their repurposing in TAMs

We wanted to repurpose existing drugs with other indications and known molecular targets to interfere with the eight selected proteins obtained via bulk and single-cell RNA-Seq and network analysis (Figure 4A). To this end, we first retrieved and prepared the 3D structures of the selected protein targets from public repositories and generated a pharmacophore model containing relevant binding pockets for each one of the protein targets (Figure 4B). We screened the pharmacophore models with 1647 FDA-approved chemical-compounds contained in the Zinc15 database [44]. We selected the drug-target interactions with a FIT score higher than one. The virtual screening resulted in 266 predictions across the eight selected target proteins (Table S3). We found 39 drugs that target at least two out of eight selected proteins (Figure 4C). Among them, six drugs have affinities towards four or more target proteins. We further filtered the virtual screening results for the largest common drug-target sets and identified four drugs (ZINC000003830943; ZINC000003830944; ZINC000003978028 (a.k.a. Clindamycin); and ZINC000008214681 (Streptomycin)) predicted to bind to four target proteins (CASP1; YBX1; ISG15; and PTGS2) (Figure 4D).

**Figure 4.**
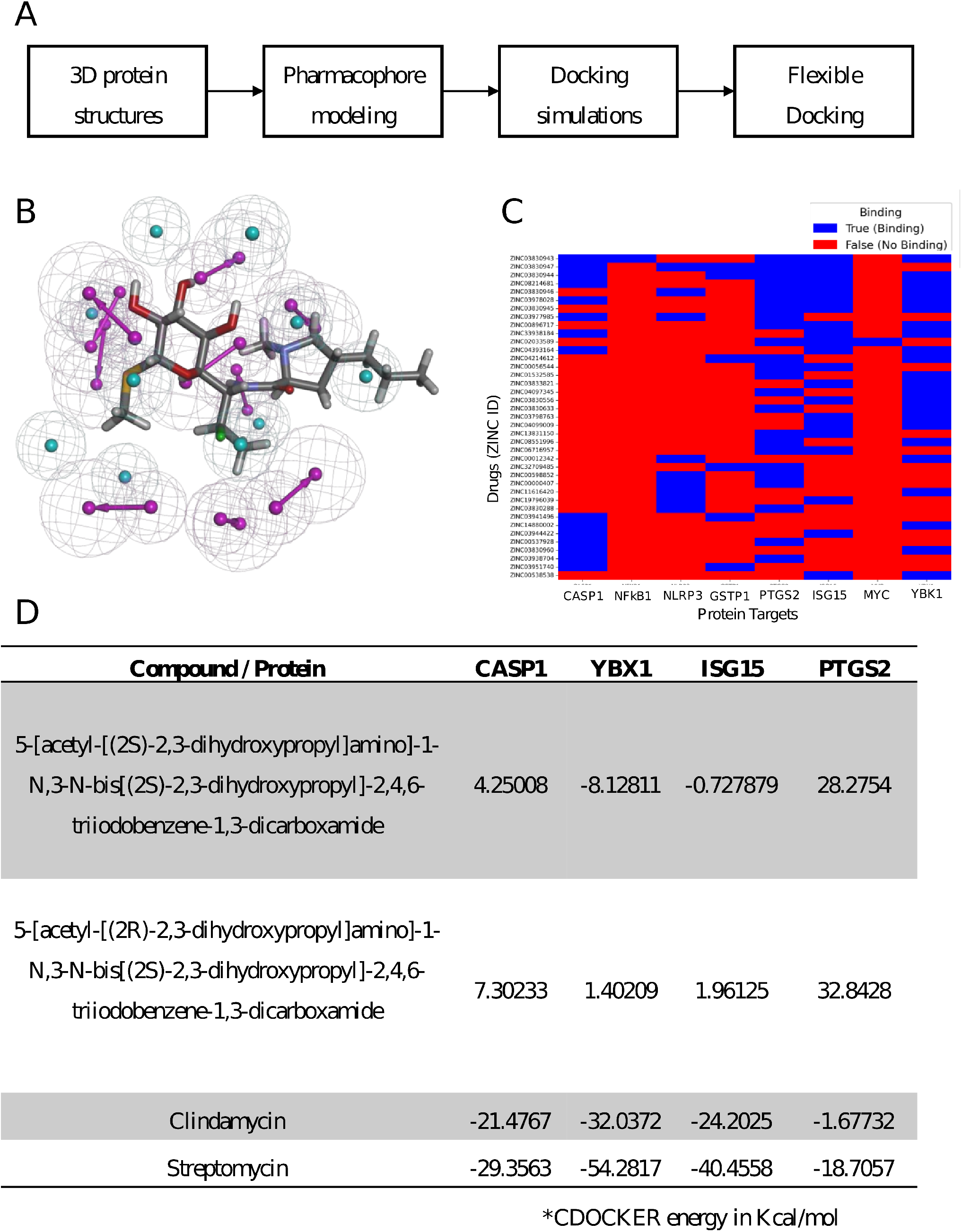
Computational screening of chemical compounds to selected protein targets. A) Central steps for the molecular docking. B) Exemplary pharmacophore of Clindamycin. C) Heatmap highlighting drug-target combinations. All drugs that have binding affinities for at least two proteins among eight selected target proteins are shown. **D) Four chemical compounds and their predicted CDOCKER energy from revised flexible docking.** CDOCKER energy unit is depicted in Kcal/mol. Only Clindamycin and Streptomycin showed suitable affinity values towards all the four target proteins.

To further confirm the predicted interactions between these drugs and the four proteins, we extracted the detailed molecular structure of the protein binding sites from the experimental literature, performed flexible docking simulations of the drugs in the binding sites, and ranked the drug-protein interactions based on their calculated CDOCKER energy (Figure 4D). To this end, we considered the top ten conformations for each of the drug–protein combinations. Our results suggest that Clindamycin and Streptomycin have better binding affinity with all the analyzed protein targets than the other two drugs considered for refined docking simulations (Figure 5). Figure 5A contains detailed information concerning the binding simulation of Clindamycin to Caspase-1, one of the most relevant protein targets linking our analysis to the TAMs immunomodulation and depletion (Table S4). Since Clindamycin has not been utilized in the context of cancer therapy and experiments to check its effect on the predicted target were considered feasible, we selected this drug for performing *in-vitro* verification experiments.

**Figure 5.**
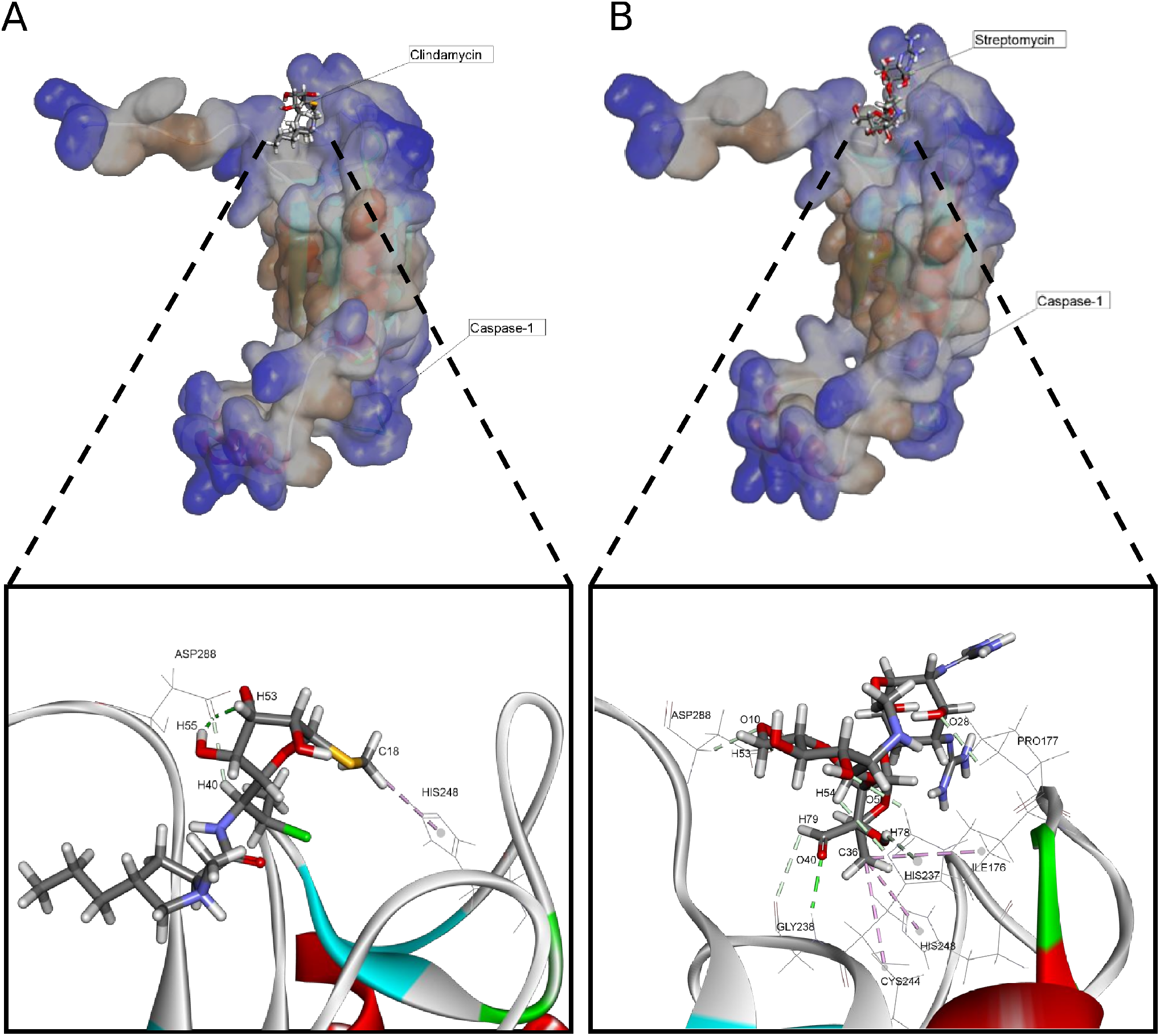
Docking simulations predicting the binding of Clindamycin (A) and Streptomycin (B) to Caspase-1. **Top:** general view of the protein and small molecule interaction**. Bottom**: detail of the binding pocket with molecular bonds highlighted.

### *In vitro* experiments confirmed that the drug-repurposing candidate Clindamycin reduces macrophage cell death via pyroptosis

Given that a) inflammasomes, and precisely NLRP3, have been implicated in solid tumor progression [51,52], b) NLRP3 is upstream the processing and activation of Caspase-1, and c) Caspase-1 activation or inhibition can be easily measured, we experimentally assessed the effect of Clindamycin on NLRP3-inflammasome activation in human macrophages (Figure 6A). To this end, we treated LPS pre-activated macrophages with Nigericin in the presence or absence of Clindamycin. Nigericin is a microbial toxin that triggers the NLRP3 inflammasome-dependent induction of IL-1β and IL-18 [53]. After treatment with Nigericin, we observed an increased amount of IL-1β (LPS: 144±61 pg/ml vs. LPS + Nigericin: 3722±748 pg/ml) and IL-18 (LPS: 17±11 pg/ml vs. LPS + Nigericin: 1084±212 pg/ml) in the supernatant after 24-hour treatment (Figure 6B). The addition of Clindamycin showed a significant reduction in Nigericin-mediated secretion of IL-1β (LPS + Nigericin: 3722±748 pg/ml vs. LPS + Nigericin + Clindamycin: 2115±961 pg/ml; p=0.006) and IL-18 (LPS + Nigericin: 1084±212 pg/ml vs. LPS + Nigericin + Clindamycin: 664±171 pg/ml; p=0.001) (Figure 6C), suggesting that Clindamycin suppresses inflammasome activity.

**Figure 6.**
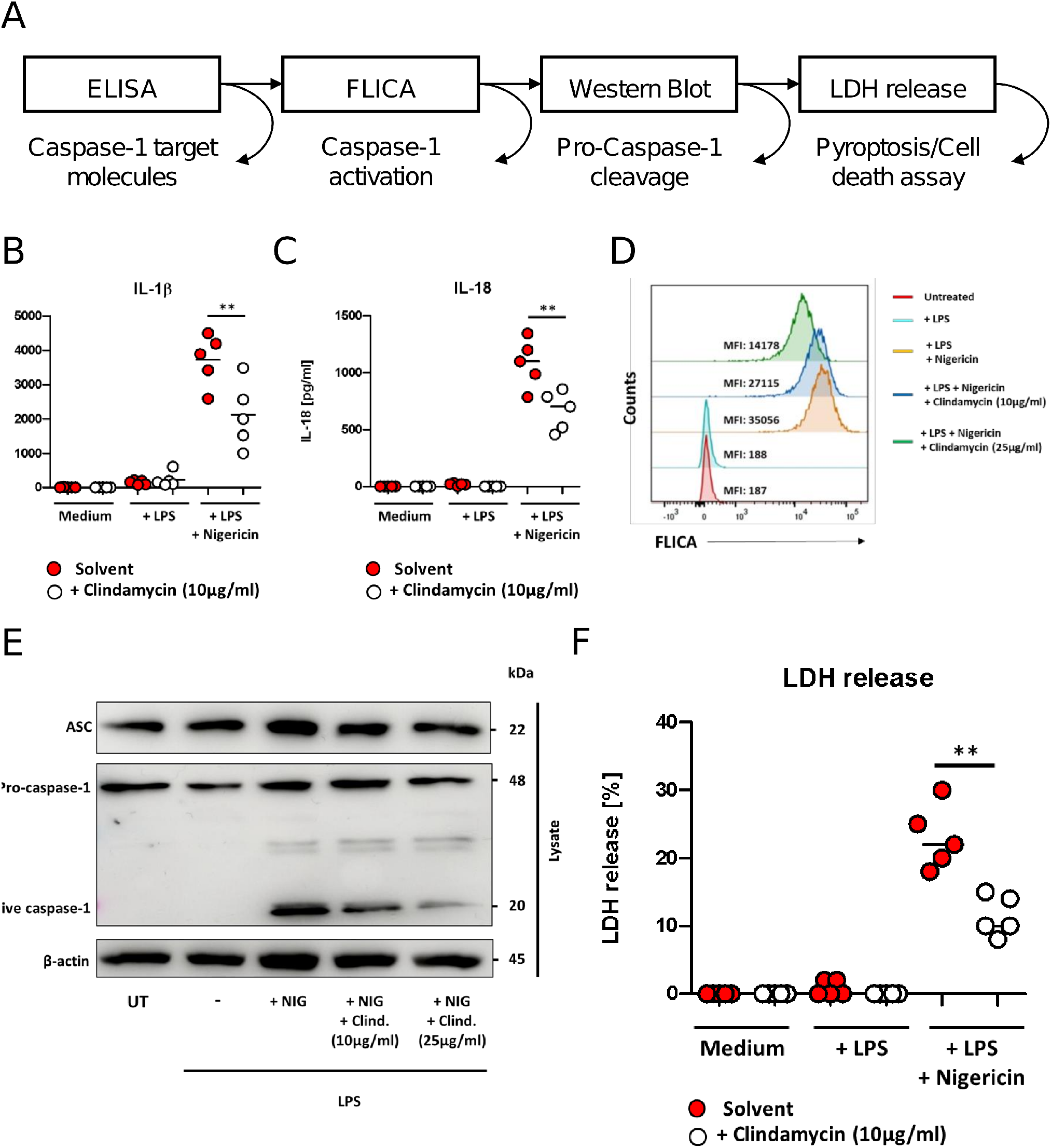
Experimental validation of the predicted interaction. A) In-vitro experiments to verify inhibition of Caspase-1 activation by Clindamycin. Monocyte derived macrophages were pretreated with (white circle) or without (red circle) Clindamycin (2h, 10µg/ml). After this, macrophages were treated with LPS (100ng/ml) alone or with LPS and Nigericin (10 µM) for 3 hours. Supernatants were analyzed by ELISA. Monocyte-derived macrophages were treated with LPS (100ng/ml, 24h) and Nigericin (10µM, 24h) in the presence or absence of Clindamycin (as indicated) and cells were analyzed for (**B**, **C**) cytokine secretion (IL1β and IL-18) by ELISA, (**D**) Caspase-1 activation by flow cytometry, (**E**) Caspase-1 activation and cleavage by Western blot or (**F**) LDH release by ELISA.

To test whether Clindamycin specifically inhibits Caspase-1, we measured Caspase-1 activation in Nigericin-treated macrophages using FLICA® reagent, a cell-permeant fluorescent-labeled inhibitor binding specifically and covalently to active Caspase-1 [52]. Flow cytometry showed an increase in Caspase-1-positive macrophages after treatment with Nigericin in comparison to only LPS-treated macrophages (Figure 6D). In contrast, we found a dose-dependent reduction in Nigericin-mediated Caspase-1 activation following pretreatment with Clindamycin (up to a 2.5-fold reduction, Figure 6D). To verify the observed inhibitory effect of Clindamycin on Caspase-1 activity, we monitored cleavage of Caspase-1 by western blot analysis. Nigericin treatment triggered cleavage of Pro-Caspase-1 to the active form of Caspase-1 (p20, 20 kDa), resulting in an additional band in the Western blot (Figure 6E). In contrast, additional pretreatment with Clindamycin reduced the fraction of active Caspase-1 in a dose-dependent manner. In the same cell lysates, we could not detect any reduction of the adapter molecule ASC (Apoptosis-associated speck-like protein containing a CARD). Given that the adapter protein ASC is upstream of the signaling pathway from Caspase-1 and binds directly to Caspase-1 for activation, this suggests that Clindamycin acts directly on Caspase-1.

Since treatment with Nigericin results in pyroptosis and Caspase-1 is a key protein for its induction, we speculated that Clindamycin may prevent the triggering of pyroptosis. To test this, we incubated macrophages with Nigericin in the presence or absence of Clindamycin, and measured the concentration of LDH, a marker of pyroptosis, in the supernatant by ELISA after 24 hours (Figure 6F). Nigericin treatment resulted in an increased release of LDH (LPS: 0.8±0.5% vs. LPS + Nigericin: 23±2%). In contrast, additional treatment with Clindamycin significantly reduced LDH release (LPS + Nigericin + Clindamycin: 11±1%; p=0.0013), suggesting that Clindamycin-mediated inhibition of Caspase-1 induces less pyroptosis in macrophages.

## Discussion

TAMs are prominent infiltrating immune cells in UM liver metastases, and their abundance positively correlates with a worse prognosis for patients [4,5]. Apart from that, TAMs are known to affect negatively ICI therapy in other cancer entities [1,2]. Since even combined ICI therapies fail to substantially increase the survival times of UM patients with liver metastases, we hypothesize we may be able to remodel the tumor microenvironment and sensitize metastases for ICI via drug-based immunomodulation of UM-associated macrophages [3]. To circumvent the time-consuming and costly *de novo* design and approval of drugs, we used computational drug repurposing to identify drug-protein interactions for immunomodulation of UM-associated macrophages.

### A scRNA-Seq data-driven network analysis identifies molecular targets for immunomodulation of TAMs

In our workflow, we first created a regulatory network involving protein and gene interactions linked to TAM-biology. We generated a network with 3863 genes and 9073 edges. To narrow the network to genes specific for UM, we combined scRNA-Seq data from uveal melanoma TAMs and healthy macrophages and isolated a small, highly-interconnected core network of 74 genes. This procedure can be further improved by using a more comprehensive collection of UM-associated macrophages and other tissues of origin for our control macrophages. Since the differential expression is only one factor for the extraction of the core network, we consider our data to be sufficient.

We combined quantitative data and expert knowledge to select the protein targets for drug repurposing. One criterion to prioritize targets was the assessment of experimental feasibility, utilized to minimize the potential laboratory work necessary to find effective drug-protein interactions. We selected eight protein targets, namely, YBX1, MYC, GSTP1, PTGS2, NLRP3, NFKB1, ISG15, and CASP1. YBX1 is a DNA and RNA binding protein whose elevated expression is linked with macrophage infiltration and poor prognosis in luminal breast cancer [54]. C-Myc (MYC) is a cell cycle and apoptosis gene known to play a pivotal role in cancer progression in multiple cancers. Pello and coworkers found that the inhibition of c-MYC in myeloid cells hampers the maturation of TAMs and impairs their pro-tumoral activity [55]. GSTP1 is a detoxifying enzyme, and its aberrant expression in breast cancer TAMs promotes IL-6 expression and drug resistance in MCF-7 in vitro experiments [56]. PTGS2 is an enzyme acting as dioxygenase or peroxidase, which participates in prostaglandin biosynthesis and inflammation. Li and coworkers found that PTGS2 is connected to the induction and maintenance of the anti-inflammatory M2 polarization in TAMs [57]. The NLRP3 inflammasome complex is an upstream activator of NF-kappaB signaling-mediated inflammatory response. Lee et al. found an association between the inhibition of the NLRP3 inflammasome in macrophages and the suppression of the metastatic potential in melanoma tumor cells [58]. ISG15 is a ubiquitin-like protein interacting with its intracellular target proteins upon activation of interferon signaling. The secretion of ISG15 by tumor cells induces an M2-like phenotype in macrophages and contributes to tumor progression and immunosuppression [59]. CASP1 is a caspase that participates in the execution phase of cell apoptosis and is involved in inflammation and cell death. Niu et al. found that Caspase-1 potentiates the pro-tumor action of TAMs [60]. Taken together, we found in the literature evidence of the connection between the selected protein targets and the activity of macrophages and TAMs.

### Pharmacophore modelling as a way of speeding up drug repurposing in TAMs

We decided to interfere with the protein targets utilizing *de novo* drug repurposing, that is, to repurpose drugs to molecular targets other than their initially approved ones. One can expand the pool for potential drugs interfering with the selected molecular targets by considering known interactions from databases like DrugBank [61]. However, repurposing to known protein-drug interactions is often biased towards well-investigated proteins. Combining both approaches could offer the possibility of considering well-known drug-target interactions for thoroughly investigated proteins and *de novo* drug repurposing for less popular ones.

We utilized pharmacophore-based analysis of an extensive database of FDA-approved drugs to identify drug-protein interactions. Pharmacophore modelling is a methodology that uses the protein active sites as potential chemical fingerprints for drug screening. This way, one can reduce the computational resources necessary to simulate the binding between the protein target and the drug, making the systematic computational screening of large libraries of active compounds possible. In our case, this resulted in 266 relevant drug-protein interactions, with four drugs being able to bind to four of the eight selected protein targets. We employed flexible docking simulations to further elaborate on the interactions between these more promising four drugs with their target proteins. This procedure is more demanding regarding computational power and manual curation but gives fine-detail predictions for the interactions. Our simulations indicate that two drugs bound significantly better to all the four targets than the others, namely Streptomycin and Clindamycin. Streptomycin is a broad-spectrum antibiotic inhibiting both Gram-positive and Gram-negative bacteria, and its described mechanism of action is the inhibition of bacteria protein synthesis. Clindamycin is an antibiotic with a bacteriostatic effect, used primarily to treat anaerobic infections, whose mechanism of action relies also on bacterial protein synthesis inhibition. Interestingly, in recent times, antibiotics have been proposed as repurposed drugs for cancer, and several clinical trials are investigating their efficacy as anticancer therapy [62]. For experimental validation, we focused on Clindamycin because it is an inexpensive compound, not been tested in the context of cancer, and has not been associated yet with Caspase-1 in the literature.

### *In vitro* tests confirm the ability of Clindamycin to interfere with the NLRP3/Caspase-1 axins in macrophages

Although our computational analysis indicates that this drug can interact with four of our top target candidates, we focus the experimental investigation on the interaction with Caspase-1 due to its central effect on inflammasome activity. Active inflammasome-carrying TAMs often promote augmented inflammation in the tumor microenvironment [52]. NLRP3, a widely studied inflammasome complex, has been directly implicated in cancer progression [51]. Activated NLRP3 recruits the adaptor molecule ASC, which binds to Pro-Caspase-1 and triggers autocatalytic activation. Active Caspase-1 catalyzes the cleavage of the pro-cytokines IL-1β and IL-18, which is necessary for their secretion and activation [63]. Further, Caspase-1 activation can trigger pyroptosis, a programmed immune-cell death characterized by plasma-membrane rupture and release of pro-inflammatory intracellular content [14]. Pyroptosis in the tumor microenvironment produces a chronic inflammatory milieu that enhances cancer cell transformation and promotes immune escape [64]. Moreover, recent findings report a blockade of IL-1β activity to be able to elicit TAM reprogramming and a decreasing inflammation [6]. Having this in mind, we hypothesized that therapeutic blockade of the TAMs inflammasome via the NLRP3/Caspase-1 axis represents a novel therapeutic strategy for the immunomodulation of TAMs, and decided to focus our drug repurposing experimental validation on this process [65]. In our experiments we found that Clindamycin indeed suppressed inflammasome activity-mediated secretion of IL-1β and IL-18 in LPS pre-activated macrophages treated with Nigericin, an NLRP3-activating microbial toxin. Our data further indicated that this effect happens downstream of ASC in the NLRP3-ASC-Caspase-1 signaling pathway, suggesting that Clindamycin acts directly on Caspase-1. Finally, we found that Clindamycin-mediated inhibition of Caspase-1 reduced pyroptosis in macrophages. We performed the experiments with macrophages derived from monocytes isolated from the peripheral blood of healthy donors due to the great difficulty of obtaining viable TAMs from UM liver metastases. However, the use of UM-associated macrophage-specific transcriptomics data to specify the core network and the presented experimental setup allows for extending the conclusions to TAMs.

With our *in-silico* approach we were able to predict a novel drug-protein interaction that proved to be immunomodulatory *in vitro*. Further preclinical *in vivo* experiments with animal models harbor the potential to solidify the inflammation-inhibiting effect of Clindamycin on macrophages in proximity to the viable tumor and could thereby uncover how this influences the susceptibility of metastatic UM to ICI.

In conclusion, we hereby propose a network-oriented methodology for *de novo* drug repurposing, which allows for filtering and prioritization of drug-target interactions. We were able to predict a new drug-target interaction that effectively blocks Caspase-1-mediated inflammasome activity *in vitro* and is therefore clinically promising for the improvement of ICI therapies for metastatic uveal melanoma. We designed the workflow having the context of TAMs in UM liver metastasis in mind, although the general methodology and its key-findings can be applied to various other implications.

## Acknowledgements

We thank Martin Eberhardt for his contribution in setting up the vcells.net.

## Contributions

JV and AW developed the concept. DC and AW performed the network reconstruction under the supervision of JV and GK. AW performed the sequencing data analysis and core network detection under the supervision of JV and XL. KPS performed the pharmacophore modelling and docking simulations under the supervision of SG. CF and JB conducted the *in vitro* experiments under the supervision of HB. JV, AW, HB, KPS, and SG drafted the manuscript. All the authors edited, corrected, and approved the submitted draft.

## Abbreviations

UM: Uveal melanoma
ICI: Immune checkpoint inhibitor
TAM: Tumor-associated macrophages
LDH: Lactate dehydrogenase
scRNA-Seq: single-cell RNA Sequencing
FC: Fold-change
TPM: Transcripts per million
BC: Betweenness centrality
D: Degree
NCBI: National Center for Biotechnology Information
GEO: Gene Expression Omnibus
HRP: Horseradish peroxidase

