## Supplementary figures and tables for "The integration of network biology and pharmacophore modeling suggests repurposing Clindamycin as an inhibitor of pyroptosis via Caspase-1 blockage in tumor-associated macrophages"

### Supplementary Material

### Affiliation

[1] Department of Dermatology, Uniklinikum Erlangen and Friedrich-Alexander Universität (FAU) Erlangen-Nürnberg, 91054 Erlangen, Germany

[2] Comprehensive Cancer Center Erlangen-European Metropolitan Area of Nuremberg (CCC ER-EMN),91054 Erlangen

[3] Deutsches Zentrum Immuntherapie (DZI),91054 Erlangen

[4] Department of Internal Medicine 5, Hematology and Oncology, Uniklinikum Erlangen, Friedrich-Alexander Universität (FAU) Erlangen-Nürnberg, 91054 Erlangen, Germany

[5] Department of Systems Biology and Bioinformatics, Universtität Rostock, Rostock, Germany

[6] Department of Internal Medicine 3, Rheumatology, Uniklinikum Erlangen, Friedrich-Alexander Universität (FAU) Erlangen-Nürnberg, 91054 Erlangen, Germany

[7] Department of Rheumatology and Clinical Immunology, Charité - Universitätsmedizin Berlin, Germany

[8] MEDI, Faculty of Medicine and Health Technology, Tampere University, Tampere, Finland

[†] Equal first authors

[$] Equal last authors

**Corresponding Author:** Laboratory of Systems Tumor Immunology, Dept. Dermatology, Univerklinikum Erlangen and Friedrich-Alexander Universität Erlangen-Nürnberg, Hartmannstr. 14, 91052, Erlangen, web: www.jveralab.net,

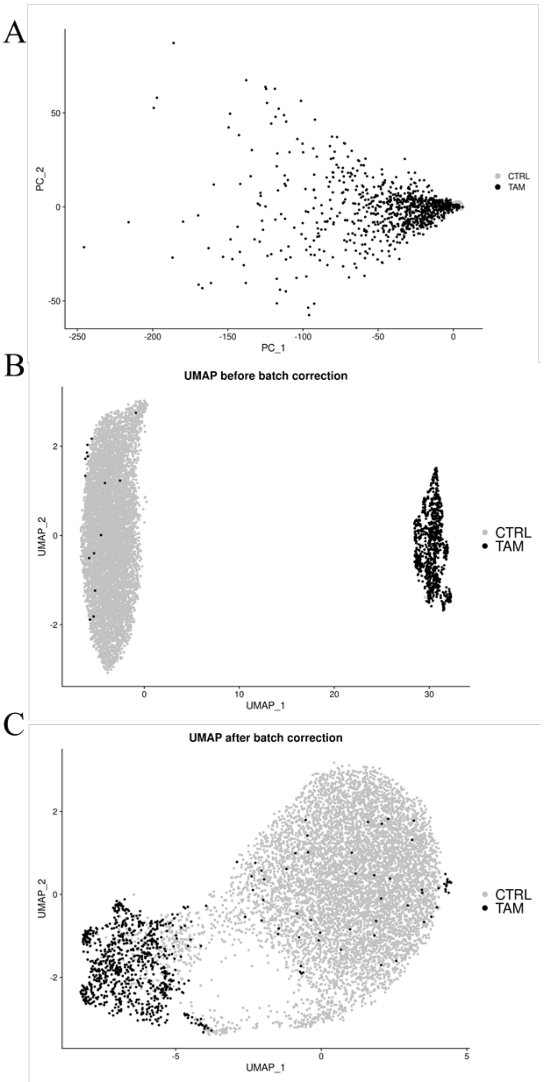


**Figure S1. Dimensionality reduction plots of combined single-cell data. A) PCA of the combined dataset.** The combined dataset shows high variance that can hardly be explained by the first two principal components. **B) UMAP of the combined dataset before batch correction.** Healthy macrophages (CTRL) and tumor-associated macrophages (TAM) show substantial batch effects and thereby form distinct clusters without any intermediate cells. **C) UMAP of the combined dataset after batch correction.** CTRL and TAM cells still form distinct clusters but several intermediate and sub-clusters emerge after correcting for batch effects with Harmony.

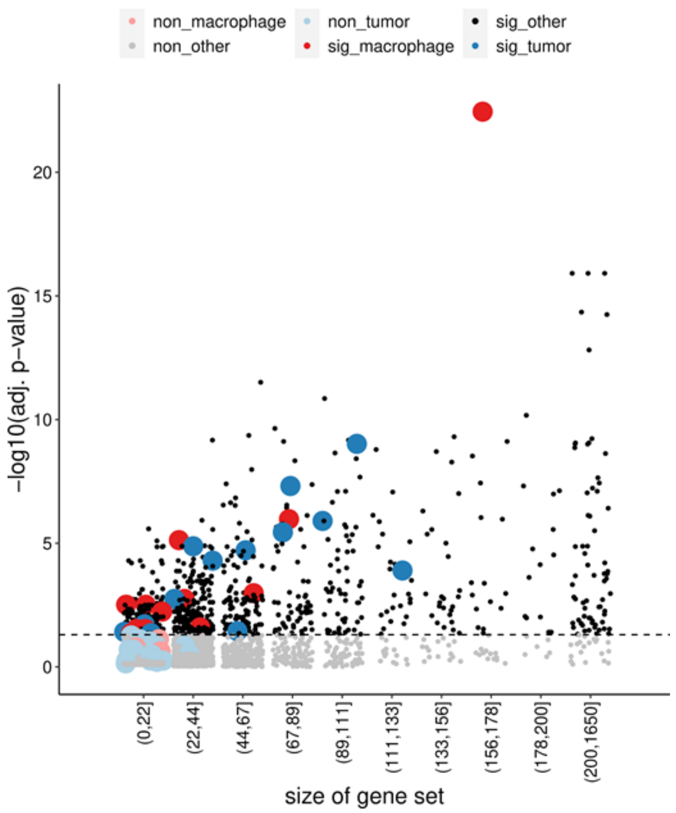


**Figure S2. Distribution of the sizes of gene sets with significantly enriched phenotypes in the GSEA.** The gene sets are classified into the corresponding groups (x-axis) based on their sizes, and the tick label is an interval representing the minimum and maximum gene sizes in a group. The y-axis is the negative value of the log10 transformation of the adjusted p-value, and the gene sets with adjusted p-values greater than 0.05 (below the dashed line) are shaded. Macrophage-related and tumor related gene sets are highlighted in red and blue, respectively.

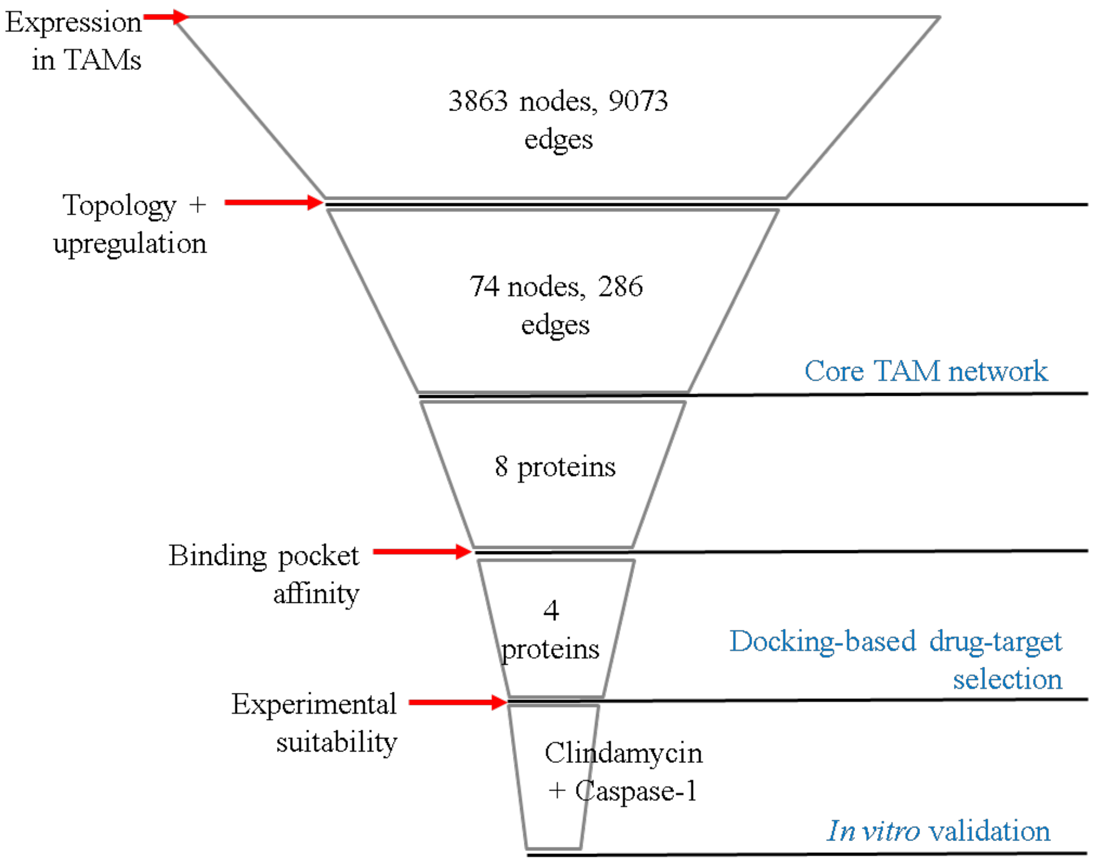


**Figure S3. Funnel plot to illustrate the step-by-step restriction to select the most promising candidate for drug repurposing experiments.** We made use of different information sources to decide which filters and restrictions to use for each of the critical steps (red arrows). We made use of different methods to solidify isolated findings (blue font).

**Supplementary Table 1. Overview of TAM network motif detection**

| Motif | Occurrences |
|---|---|
| 2-nodes-2-edges Feedback Loop | 30 |
| 3-nodes-3-edges Feedback Loop | 25 |
| 3-nodes-4-edges Feedback Loop | 14 |
| 4-nodes-4-edges Feedback Loop | 65 |
| 3-nodes-3-edges Forward Loop | 2990 |
| 3-nodes-4-edges Forward Loop | 270 |
| 4-nodes-4-edges Forward Loop | 5641 |

### Supplementary Table 2. Target selection based on core network and expression data

| HGNC | Log2FC | BC | EC | normLog2FC | normBC | normEC | Score |
|---|---|---|---|---|---|---|---|
| YBX1 | 1.41 | 7.0375 | 49 | 0.306 | 0.644 | 1 | 1.95 |
| MYC | 0.35 | 10.9277 | 30 | 0.016 | 1 | 0.604 | 1.62 |
| STAT1 | 2.68 | 0.7598 | 17 | 0.655 | 0.06953 | 0.333 | 1.058 |
| SPARC | -3.94 | 0 | 3 | 1 | 0 | 0.042 | 1.042 |
| HIF1A | 1.46 | 3.0593 | 21 | 0.319 | 0.27995 | 0.417 | 1.016 |
| GSTP1 | 3.61 | 0 | 4 | 0.91 | 0 | 0.062 | 0.972 |
| PTGS2 | 2.87 | 0 | 10 | 0.707 | 0 | 0.188 | 0.895 |
| IRF1 | 1.33 | 1.4761 | 21 | 0.284 | 0.13508 | 0.417 | 0.836 |
| PNRC1 | 2.88 | 0 | 2 | 0.708 | 0 | 0.021 | 0.729 |
| NLRP3 | 2.48 | 0 | 4 | 0.6 | 0 | 0.062 | 0.663 |
| STAT4 | 0 | 5.4158 | 9 | 0 | 0.4956 | 0.167 | 0.662 |
| ALOX5AP | -2.43 | 0 | 4 | 0.586 | 0 | 0.062 | 0.648 |
| REL | 2.36 | 0 | 4 | 0.566 | 0 | 0.062 | 0.628 |
| RELA | 0 | 2.566 | 18 | 0 | 0.23482 | 0.354 | 0.589 |
| NFKB1 | 0.73 | 1.1894 | 17 | 0.119 | 0.10884 | 0.333 | 0.561 |
| AKT1 | -0.72 | 1.8747 | 14 | 0.116 | 0.17155 | 0.271 | 0.558 |
| CXCR4 | 2.02 | 0 | 5 | 0.474 | 0 | 0.083 | 0.557 |
| ISG15 | 2.14 | 0 | 3 | 0.507 | 0 | 0.042 | 0.549 |
| XRCC5 | 1.86 | 0.3548 | 3 | 0.429 | 0.03247 | 0.042 | 0.503 |
| ZFP36 | 1.76 | 0 | 5 | 0.402 | 0 | 0.083 | 0.485 |
| JUN | 0 | 1.3428 | 18 | 0 | 0.12288 | 0.354 | 0.477 |
| STAT3 | 0.42 | 0.5765 | 19 | 0.034 | 0.05276 | 0.375 | 0.461 |
| ADAR | 1.4 | 0.8758 | 4 | 0.303 | 0.08014 | 0.062 | 0.446 |
| PRKACB | 0 | 1.8418 | 14 | 0 | 0.16854 | 0.271 | 0.439 |
| NFKBIA | 1.52 | 0 | 5 | 0.337 | 0 | 0.083 | 0.42 |
| ETS2 | 1.32 | 0.1118 | 7 | 0.281 | 0.01023 | 0.125 | 0.416 |
| NR4A1 | 1.62 | 0 | 2 | 0.364 | 0 | 0.021 | 0.385 |
| MAPK1 | -0.29 | 1.8275 | 11 | 0 | 0.16723 | 0.208 | 0.376 |
| FOS | 0.71 | 0.0594 | 13 | 0.115 | 0.00544 | 0.25 | 0.37 |
| PRKCD | -1.1 | 0.1716 | 7 | 0.22 | 0.0157 | 0.125 | 0.361 |
| MAPK9 | 0 | 2.2664 | 8 | 0 | 0.2074 | 0.146 | 0.353 |
| CCND1 | 0.8 | 0 | 9 | 0.138 | 0 | 0.167 | 0.305 |
| HMOX1 | 0.4 | 1.0901 | 9 | 0.03 | 0.09976 | 0.167 | 0.297 |
| CHUK | 0 | 1.2332 | 9 | 0 | 0.11285 | 0.167 | 0.28 |
| RUNX1 | -0.57 | 0.9608 | 6 | 0.076 | 0.08793 | 0.104 | 0.269 |
| PIK3CA | 0 | 0.828 | 10 | 0 | 0.07577 | 0.188 | 0.263 |
| SREBF1 | 0 | 0.0817 | 12 | 0 | 0.00748 | 0.229 | 0.237 |
| EPAS1 | 0.58 | 0 | 8 | 0.08 | 0 | 0.146 | 0.226 |
| CXCL2 | 0.57 | 0 | 7 | 0.075 | 0 | 0.125 | 0.2 |
| IKBKB | 0 | 0.1298 | 10 | 0 | 0.01187 | 0.188 | 0.199 |
| ATF2 | 0 | 0.1049 | 10 | 0 | 0.0096 | 0.188 | 0.197 |
| ATF1 | -0.43 | 0 | 8 | 0.039 | 0 | 0.146 | 0.184 |
| CASP8 | -0.83 | 0 | 2 | 0.147 | 0 | 0.021 | 0.168 |
| CIITA | 0.75 | 0 | 3 | 0.124 | 0 | 0.042 | 0.166 |
| PRKCA | 0.48 | 0.094 | 6 | 0.051 | 0.00861 | 0.104 | 0.163 |
| MAP2K1 | -0.53 | 0.2021 | 4 | 0.066 | 0.0185 | 0.062 | 0.147 |
| CDKN1A | 0 | 0 | 8 | 0 | 0 | 0.146 | 0.146 |
| ACLY | -0.59 | 0 | 4 | 0.081 | 0 | 0.062 | 0.143 |
| MAPKAPK2 | -0.5 | 0.0175 | 5 | 0.056 | 0.0016 | 0.083 | 0.14 |
| PTPRE | 0.63 | 0.2021 | 2 | 0.093 | 0.0185 | 0.021 | 0.133 |
| STAT6 | 0 | 0.1298 | 6 | 0 | 0.01188 | 0.104 | 0.116 |
| MAP3K7 | 0 | 0.0715 | 6 | 0 | 0.00655 | 0.104 | 0.111 |
| SOS1 | 0.61 | 0 | 2 | 0.086 | 0 | 0.021 | 0.107 |
| CASP1 | 0.44 | 0.0168 | 4 | 0.041 | 0.00154 | 0.062 | 0.105 |
| ITPR3 | 0 | 0 | 6 | 0 | 0 | 0.104 | 0.104 |
| RPS6KB1 | -0.52 | 0.1409 | 2 | 0.062 | 0.01289 | 0.021 | 0.096 |
| RPS6KA1 | 0 | 0.1015 | 5 | 0 | 0.00928 | 0.083 | 0.093 |
| MAP2K6 | 0 | 0.006 | 5 | 0 | 0.00055 | 0.083 | 0.084 |
| ADCY1 | 0 | 0.0042 | 5 | 0 | 0.00038 | 0.083 | 0.084 |
| EGLN3 | 0 | 0.0028 | 5 | 0 | 0.00025 | 0.083 | 0.084 |
| PRKCG | 0 | 0 | 5 | 0 | 0 | 0.083 | 0.083 |
| PDPK1 | 0 | 0 | 5 | 0 | 0 | 0.083 | 0.083 |
| SUMO3 | -0.51 | 0 | 2 | 0.06 | 0 | 0.021 | 0.081 |
| CXCL2 | 0.57 | 0 | 1 | 0.075 | 0 | 0 | 0.075 |
| PRKCZ | 0 | 0.0715 | 4 | 0 | 0.00655 | 0.062 | 0.069 |
| LTA4H | 0.46 | 0 | 2 | 0.046 | 0 | 0.021 | 0.067 |
| MAP3K14 | 0 | 0 | 4 | 0 | 0 | 0.062 | 0.062 |
| NLRP1 | 0 | 0 | 4 | 0 | 0 | 0.062 | 0.062 |
| INPP5D | -0.34 | 0.0523 | 3 | 0.014 | 0.00479 | 0.042 | 0.06 |
| CASP7 | 0 | 0 | 3 | 0 | 0 | 0.042 | 0.042 |
| PIAS3 | 0 | 0.2054 | 2 | 0 | 0.01879 | 0.021 | 0.04 |
| THEM4 | 0 | 0 | 2 | 0 | 0 | 0.021 | 0.021 |

| | | | | | | | |
|---|---|---|---|---|---|---|---|
| ODC1 | 0 | 0 | 2 | 0 | 0 | 0.021 | 0.021 |
| VHL | 0 | 0 | 2 | 0 | 0 | 0.021 | 0.021 |
| CREBBP | 0 | 0 | 2 | 0 | 0 | 0.021 | 0.021 |

**Supplementary Table 3. Virtual screening results (4/8)**

YBX1

| Name | Fit value |
|---|---|
| ZINC00005626 | 4,39888 |
| ZINC03831401 | 4,01592 |
| ZINC08551996 | 3,63188 |
| ZINC03978028 | 3,54797 |
| ZINC04544045 | 3,26216 |
| ZINC08551995 | 3,05089 |
| ZINC00621853 | 3,04217 |
| ZINC33938184 | 3,01327 |
| ZINC03812841 | 2,80564 |
| ZINC08214681 | 2,73719 |
| ZINC03833846 | 2,55174 |
| ZINC04099009 | 2,50754 |
| ZINC04393164 | 2,50342 |
| ZINC03830996 | 2,50338 |
| ZINC03830335 | 2,49607 |
| ZINC08214681 | 2,49419 |
| ZINC03830581 | 2,39051 |
| ZINC03831269 | 2,37388 |
| ZINC04097345 | 2,37198 |
| ZINC12503222 | 2,30447 |
| ZINC03831357 | 2,24644 |
| ZINC01552908 | 2,09155 |
| ZINC00119988 | 2,00827 |
| ZINC03830709 | 2,00654 |
| ZINC03833821 | 1,845 |
| ZINC01532585 | 1,83496 |
| ZINC04097345 | 1,72299 |
| ZINC08214681 | 1,69527 |
| ZINC14880002 | 1,68232 |
| ZINC03830943 | 1,67349 |
| ZINC03830633 | 1,6461 |
| ZINC11592735 | 1,61924 |
| ZINC04215367 | 1,61154 |
| ZINC06920406 | 1,60782 |
| ZINC03830556 | 1,58266 |
| ZINC03830944 | 1,52107 |
| ZINC04214612 | 1,42348 |
| ZINC01846741 | 1,41397 |
| ZINC00538538 | 1,38925 |
| ZINC03830945 | 1,37778 |
| ZINC00643114 | 1,3648 |
| ZINC03830961 | 1,36135 |
| ZINC03798763 | 1,34227 |
| ZINC03830944 | 1,10767 |
| ZINC03830943 | 1,07072 |
| ZINC00012342 | 1,06816 |
| ZINC03830943 | 1,0128 |

MYC

| Name | Fit value |
|---|---|
| ZINC02033589 | 2,98714 |

ISG15

| Name | Fit value |
|---|---|
| ZINC13831151 | 4,27125 |
| ZINC03830674 | 3,79187 |
| ZINC08214681 | 3,41641 |
| ZINC00057235 | 3,39196 |
| ZINC28240499 | 3,36103 |
| ZINC13831150 | 3,31277 |
| ZINC04544044 | 3,2658 |
| ZINC12484926 | 3,21323 |
| ZINC03831268 | 3,20549 |
| ZINC03830947 | 3,19947 |
| ZINC01530866 | 3,06066 |
| ZINC03831358 | 3,05275 |
| ZINC03831448 | 2,99026 |
| ZINC02033589 | 2,97596 |
| ZINC00896717 | 2,96336 |
| ZINC03830947 | 2,94613 |
| ZINC08552291 | 2,91586 |
| ZINC03830943 | 2,84096 |
| ZINC03830993 | 2,79604 |
| ZINC03830946 | 2,76688 |
| ZINC03831358 | 2,73679 |
| ZINC33938184 | 2,7181 |
| ZINC03833821 | 2,68341 |
| ZINC03978028 | 2,63298 |
| ZINC03831268 | 2,63115 |
| ZINC06716957 | 2,61439 |
| ZINC19364219 | 2,5703 |
| ZINC03964325 | 2,53902 |
| ZINC33938184 | 2,51389 |
| ZINC03831461 | 2,50915 |
| ZINC00020230 | 2,50128 |
| ZINC03794711 | 2,48776 |
| ZINC03830944 | 2,47209 |
| ZINC03875332 | 2,40774 |
| ZINC19802035 | 2,36677 |
| ZINC03830556 | 2,308 |
| ZINC08214681 | 2,306 |
| ZINC03830923 | 2,24712 |
| ZINC03977994 | 2,24553 |
| ZINC11592789 | 2,2333 |
| ZINC04393164 | 2,22656 |
| ZINC03830943 | 2,19822 |
| ZINC03798763 | 2,16557 |
| ZINC03830944 | 2,15887 |
| ZINC03830946 | 2,15482 |
| ZINC00403079 | 2,09135 |
| ZINC03830945 | 1,9635 |
| ZINC13911941 | 1,93916 |
| ZINC00056544 | 1,9253 |
| ZINC08214681 | 1,8501 |
| ZINC01542901 | 1,83934 |
| ZINC03830580 | 1,83443 |
| ZINC14879985 | 1,82305 |
| ZINC03830944 | 1,81918 |
| ZINC04099009 | 1,61177 |
| ZINC03944422 | 1,59627 |
| ZINC03978028 | 1,3794 |
| ZINC19796039 | 1,34846 |
| ZINC03830946 | 1,31236 |
| ZINC14879987 | 1,29297 |
| ZINC04097283 | 1,15402 |
| ZINC00538538 | 1,09767 |

PTGS2

| Name | Fit value |
|---|---|
| ZINC00598852 | 3,65842 |
| ZINC32709485 | 3,64234 |
| ZINC03830959 | 3,4318 |
| ZINC00057255 | 3,30341 |
| ZINC00056544 | 3,1965 |
| ZINC00537928 | 3,01043 |
| ZINC06716957 | 2,98216 |
| ZINC08551996 | 2,86545 |
| ZINC03982483 | 2,85035 |
| ZINC03830246 | 2,8356 |
| ZINC03824921 | 2,77779 |
| ZINC13831150 | 2,77144 |
| ZINC00598852 | 2,76676 |
| ZINC32709485 | 2,76592 |
| ZINC03978028 | 2,75611 |
| ZINC03977985 | 2,73803 |
| ZINC05844792 | 2,68222 |
| ZINC04097285 | 2,64679 |
| ZINC19796041 | 2,62384 |
| ZINC33956087 | 2,61227 |
| ZINC28248877 | 2,50688 |
| ZINC03830946 | 2,48577 |
| ZINC13831150 | 2,4824 |
| ZINC00601249 | 2,43958 |
| ZINC03831403 | 2,41607 |
| ZINC03830633 | 2,39177 |
| ZINC04544043 | 2,38331 |
| ZINC08143614 | 2,37702 |
| ZINC08214681 | 2,37208 |
| ZINC03830958 | 2,37006 |
| ZINC00896717 | 2,34696 |
| ZINC03830943 | 2,34102 |
| ZINC04214612 | 2,33182 |
| ZINC00000407 | 2,31226 |
| ZINC03830288 | 2,235 |
| ZINC03830947 | 2,23406 |
| ZINC32709485 | 2,1761 |
| ZINC00896512 | 2,10924 |
| ZINC04097345 | 2,09877 |
| ZINC11592620 | 2,09094 |
| ZINC00896717 | 2,07586 |
| ZINC03830947 | 2,05365 |
| ZINC03938704 | 1,88433 |
| ZINC03830947 | 1,80464 |
| ZINC01542393 | 1,661 |
| ZINC00896838 | 1,6198 |
| ZINC08214681 | 1,61095 |
| ZINC08551108 | 1,60252 |
| ZINC03794794 | 1,54986 |
| ZINC08214681 | 1,5476 |
| ZINC08552293 | 1,50677 |
| ZINC02033589 | 1,38547 |
| ZINC03830944 | 1,2727 |
| ZINC06716957 | 1,19921 |
| ZINC03830945 | 1,10662 |
| ZINC01532585 | 1,06051 |

**Supplementary Table 3. Virtual screening results (4/8)**

GSTP1

| Name | Fit value |
|---|---|
| ZINC03812892 | 3,34057 |
| ZINC01562698 | 2,61107 |
| ZINC03831365 | 2,53807 |
| ZINC32709485 | 2,46136 |
| ZINC03916973 | 2,37496 |
| ZINC03941496 | 1,9713 |
| ZINC03830947 | 1,89015 |
| ZINC04214612 | 1,65793 |
| ZINC03831449 | 1,38708 |
| ZINC03951740 | 1,35345 |
| ZINC03830944 | 1,3275 |

NLRP3

| Name | Fit value |
|---|---|
| ZINC01530862 | 4,21051 |
| ZINC00020242 | 4,04282 |
| ZINC03831296 | 3,83312 |
| ZINC03830288 | 3,76024 |
| ZINC04097451 | 3,75675 |
| ZINC11616423 | 3,7241 |
| ZINC04097451 | 3,71788 |
| ZINC19796039 | 3,63321 |
| ZINC03830381 | 3,62321 |
| ZINC00001127 | 3,47147 |
| ZINC03831089 | 3,4387 |
| ZINC03830683 | 3,4076 |
| ZINC03978005 | 3,35093 |
| ZINC03831271 | 3,18993 |
| ZINC38664623 | 3,17541 |
| ZINC26573040 | 3,10914 |
| ZINC11616420 | 3,07772 |
| ZINC00000401 | 3,07247 |
| ZINC11616423 | 3,05715 |
| ZINC03830946 | 3,00196 |
| ZINC03831271 | 2,99731 |
| ZINC04097451 | 2,98732 |
| ZINC03830285 | 2,94026 |
| ZINC00000401 | 2,88348 |
| ZINC00402980 | 2,80171 |
| ZINC16343331 | 2,78232 |
| ZINC00001127 | 2,75809 |
| ZINC03831271 | 2,73797 |
| ZINC00012342 | 2,70496 |
| ZINC00402980 | 2,70413 |
| ZINC03831271 | 2,69613 |
| ZINC03830583 | 2,68632 |
| ZINC16343331 | 2,68368 |
| ZINC05735567 | 2,59363 |
| ZINC03977985 | 2,5595 |
| ZINC16343331 | 2,5369 |
| ZINC14243816 | 2,50994 |
| ZINC03830629 | 2,45955 |
| ZINC00000401 | 2,38449 |
| ZINC00402980 | 2,37034 |
| ZINC03830683 | 2,30375 |
| ZINC00402980 | 2,28445 |
| ZINC03812923 | 2,25546 |
| ZINC00402979 | 2,24845 |
| ZINC03830947 | 2,23377 |
| ZINC03831426 | 2,138 |
| ZINC19632692 | 2,0941 |
| ZINC22921371 | 2,08814 |
| ZINC00000407 | 2,05343 |
| ZINC03831047 | 1,97706 |
| ZINC13513943 | 1,96888 |
| ZINC00598852 | 1,9102 |
| ZINC03830984 | 1,86329 |
| ZINC00009073 | 1,81718 |
| ZINC03815418 | 1,81273 |
| ZINC03815418 | 1,72291 |
| ZINC00000401 | 1,6018 |
| ZINC01530599 | 1,5285 |
| ZINC03831271 | 1,45061 |
| ZINC03831422 | 1,35836 |
| ZINC01530652 | 1,11832 |

NFKB1

| Name | Fit value |
|---|---|
| ZINC03785268 | 1,78821 |
| ZINC03830943 | 1,31651 |

CASP1

| Name | Fit value |
|---|---|
| ZINC11592627 | 3,04002 |
| ZINC08214681 | 3,03217 |
| ZINC03977985 | 3,02441 |
| ZINC03920719 | 2,58452 |
| ZINC03938746 | 2,51461 |
| ZINC03951740 | 2,46002 |
| ZINC03938704 | 2,44251 |
| ZINC03830960 | 2,36073 |
| ZINC03951740 | 2,1781 |
| ZINC00537928 | 2,14683 |
| ZINC01692922 | 2,03591 |
| ZINC03830943 | 1,80151 |
| ZINC03830947 | 1,73256 |
| ZINC03978028 | 1,71975 |
| ZINC03914596 | 1,71817 |
| ZINC03830944 | 1,71031 |
| ZINC03830947 | 1,64645 |
| ZINC03914596 | 1,58157 |
| ZINC03944422 | 1,43757 |
| ZINC14880002 | 1,33786 |
| ZINC04393164 | 1,25419 |
| ZINC03941496 | 1,2254 |
| ZINC11592628 | 1,04467 |
| ZINC33938184 | 1,01088 |

**Supplementary Table 3. Bond information of Clindamycin and Streptomycin with CASP1**

| Name | Distance | Category | Types |
|---|---|---|---|
| Clindamycin:H53 - CASP1:ASP288:OD1 | 2.21746 | Hydrogen Bond | Conventional Hydrogen Bond |
| Clindamycin:H55 - CASP1:ASP288:OD1 | 2.06199 | Hydrogen Bond | Conventional Hydrogen Bond |
| Clindamycin:H40 - CASP1:ASP288:OD1 | 2.6524 | Hydrogen Bond | Carbon Hydrogen Bond |
| CASP1:HIS248 - Clindamycin:C18 | 5.3007 | Hydrophobic | Pi-Alkyl |

| Name | Distance | Category | Types |
|---|---|---|---|
| Streptomycin:H53 - CASP1:ASP288:OD1 | 2.30275 | Hydrogen Bond | Conventional Hydrogen Bond |
| CASP1:GLY238:HN - Streptomycin:O40 | 2.29409 | Hydrogen Bond | Conventional Hydrogen Bond |
| Streptomycin:H54- CASP1:HIS237:ND1 | 3.05416 | Hydrogen Bond | Carbon Hydrogen Bond |
| Streptomycin:H79 - CASP1:GLY238:O | 3.09566 | Hydrogen Bond | Carbon Hydrogen Bond |
| CASP1:PRO177:HD1- Streptomycin:O28 | 2.86354 | Hydrogen Bond | Carbon Hydrogen Bond |
| CASP1:HIS237:HE1- Streptomycin:O5 | 2.52436 | Hydrogen Bond | Carbon Hydrogen Bond |
| CASP1:ASP288:HA - Streptomycin:O10 | 2.62191 | Hydrogen Bond | Carbon Hydrogen Bond |
| Streptomycin:H78 - CASP1:HIS237 | 2.98247 | Hydrogen Bond | Pi-Donor Hydrogen Bond |
| Streptomycin:C36 - CASP1:ILE176 | 5.17239 | Hydrophobic | Alkyl |
| Streptomycin:C36 - CASP1:CYS244 | 4.75277 | Hydrophobic | Alkyl |
| CASP1:HIS248 - Streptomycin:C36 | 4.78125 | Hydrophobic | Pi-Alkyl |
